# Urine-based detection of biomarkers indicative of chronic kidney disease in a patient cohort from Ghana

**DOI:** 10.1101/2022.10.27.513996

**Authors:** Wasco Wruck, Vincent Boima, Lars Erichsen, Chantelle Thimm, Theresa Koranteng, Edward Kwakyi, Sampson Antwi, Dwomoa Adu, James Adjaye

## Abstract

Chronic kidney disease (CKD) is a global health burden with a continuously increasing prevalence associated with an increasing incidence of diabetes and hypertension in aging populations. The CKD definition of a more than three months lasting low glomerular filtration rate (GFR) or other renal impairments including proteinuria implies that multiple factors may contribute to the disease. While there are indications of ethnic differences it is hard to disentangle these from confounding social factors. Usually, CKD is detected in later stages of the disease when irreversible renal damage has already occurred, thus suggesting a need for early non-invasive diagnostic markers.

In this study, we explored the urine secretome of a CKD patient cohort from Ghana employing a kidney-injury and a more general cytokine assay.

We identified panels of kidney-specific cytokine markers which were also gender-specific and a panel of gender-independent cytokine markers. The gender-specific markers are IL10 and MME for male and CLU, RETN, AGER, EGFR and VEGFA for female. The gender-independent cytokine markers were APOA1, ANGPT2, C5, CFD, GH1, ICAM1, IGFBP2, IL8, KLK4, MMP9 and SPP1 (up-regulated) and FLT3LG, CSF1, PDGFA, RETN and VEGFA (down-regulated).

APOA1 – the major component of HDL particles – was up-regulated in Ghanaian CKD patients and its co-occurrence with APOL1 in a subpopulation of HDL particles may point to specific CKD-predisposing APOL1 haplotypes in patients of African descent – this however needs further investigation. The identified panels may lay down the foundation for CKD-biomarker assays to be confirmed in further studies with a larger cohort of patients.

## Introduction

Chronic kidney disease (CKD) is defined by a glomerular filtration rate (GFR) <60 mL/min per 1·73 m^2^ or the presence of kidney damage predominantly manifested by proteinuria for 3 months or more ^1,2^. Protein in the urine as an indicator of kidney damage is often measured by the urine albumin-to-creatinine ratio (UACR). Proteinuria and decreased GFR directly reflect physical properties of the filter between blood and urine constituted by an endothelial layer, the glomerular basement membrane (GMB) and podocytes. This filter is coarse at the endothelial side and fine at the podocytes and works in the dimension of nanometers in addition to a negative polarity. Thus, bigger and negatively charged molecules such as most proteins under physiological conditions cannot traverse this barrier. In proteinuria, larger proteins such as albumin, immunoglobulins G and M and α_1_-microglobulin, β_2_-microglobulin, correlating with the severity of histologic lesions ^3^, can traverse. These proteins can as a consequence impair the re-absorption of other smaller molecules by the proximal tubular epithelial cells and in final stages lead to their toxic damage ^3^.

The rising prevalence of CKD is largely influenced by the increase of diabetes and hypertension among the aging population ^4,5^. Although substantial percentages of CKD patients will progress to more severe disease stages requiring dialysis or transplantation most patients die of associated cardiovascular disease (CVD) than of end stage renal disease (ESRD) ^6^. Although diabetes is the most common cause of CKD it is not clear why only 30% of patients with type 1 and 25 to 40% of patients with type 2 diabetes progress to nephropathy irrespective of glycemic control ^4, 7, 8^.

Ethnic differences have been reported for diabetic nephropathy particularly in Pima Indians ^4^ and for focal segemental glomerulosclerosis (FSGS) in patients of African descent who have higher frequencies of the FSGS-predisposing *APOL1* G1 and G2 haplotypes which on the other hand have advantages against the sleeping-sickness causing parasite-*Trypanosoma brucei brucei* ^9^.

Biomarkers for the early diagnosis of CKD are urgently needed and studies aiming at their identification have been performed using serum ^10^ as well as urine samples ^11, 12^.

In this study aiming at CKD biomarker identification, we investigated proteins secreted into the urine of a cohort of CKD patients and healthy controls from Ghana using a general (Human XL) assay and a kidney-injury specific cytokine assay.

## Results

### Patient characteristics

Urine samples from CKD patients aged 6-71 years and control urine from probands with no history of CKD aged 11-68 were used. The Urine samples were collected in Ghana at the University of Ghana Medical School. Decisive criteria for the choice of CKD urine samples were the glomerular filtration rate (GFR) <60 mL/min per 1-73 m^2^, which is documented for CKD ^13^. Furthermore CKD is associated with an elevated albumin creatine ratio (ACR) <300 mg/g ^13^. In experiment series 1 urine samples of patients from group 1 were investigated, in experiment series 2 urine samples of patients from group 2 were investigated and in experiment series 3 urine samples of patients from group 1 were investigated.

### Strategy for the identification of CKD markers

Figure 1 shows the workflow used in this study to identify CKD biomarkers. In the first phase, experiment series 1 and 2 were performed on the kidney-injury cytokine assay platform. Each experiment series consisted of 10 pooled male CKD urine samples, 10 pooled female CKD urine samples, 10 pooled male healthy control urine samples, 10 pooled female healthy control urine samples. Series 1 and 2 differed by the selection of distinct individuals. In the second phase, experiment series 3 was performed on the Human cytokine XL assay platform. The experiment series consisted of 10 pooled male CKD urine samples, 10 pooled female CKD urine samples and one male healthy control sample.

**Figure 1:**
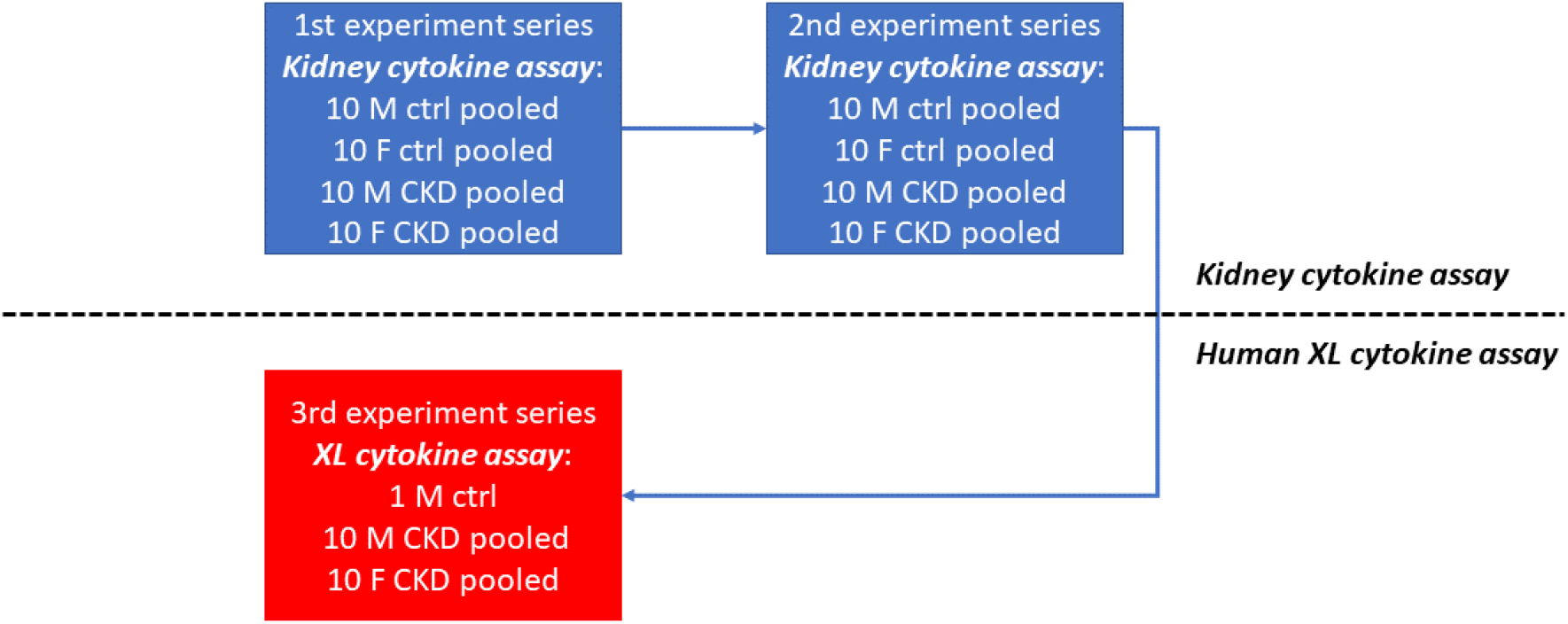
Workflow of the CKD biomarker identification. The pipeline for identification of CKD biomarkers employed the Human Kidney Biomarker Array in phase 1 of experiment series1 and 2 and the human XL cytokine assay in phase 2 with experiment series 3. Pools of 10 male and 10 female urine samples were used for CKD patients and healthy controls.

### Identification of CKD biomarkers using the Human Kidney Biomarker Array

Figure 2 shows results of experiment series 1 on the kidney cytokine assay and the identified CKD biomarkers. At the global level of the kidney cytokine expression, the data seperated (Figure 2A) into two distinct clusters of CKD (male and female) and healthy control (male and female). The heatmap (Figure 2B) and barplot (Figure 2C) show biomarkers which were found up-regulated (up) in male (M): CLU, CXCL1, IL1RN, IL10, RBP4, SKP1, down-regulated (down): DPP4, EGFR, MME, MMP9, VEGF. The heatmap (Figure 2D) and barplot (Figure 2E) show biomarkers which were found up-regulated (up) in female (F): ANPEP, ANXA5, B2M, CCL2, CCN1, CLU, CXCL16, DPP4, EGF, IL6, IL10, HAVCR1, KLK3, LCN2, MMP9, PLAU, RETN, SERPINA3, SKP1, TNFA, TNFSRF14, TTF3, down-regulated (down): AGER, EGFR, FABP1, VEGF.

**Figure 2:**
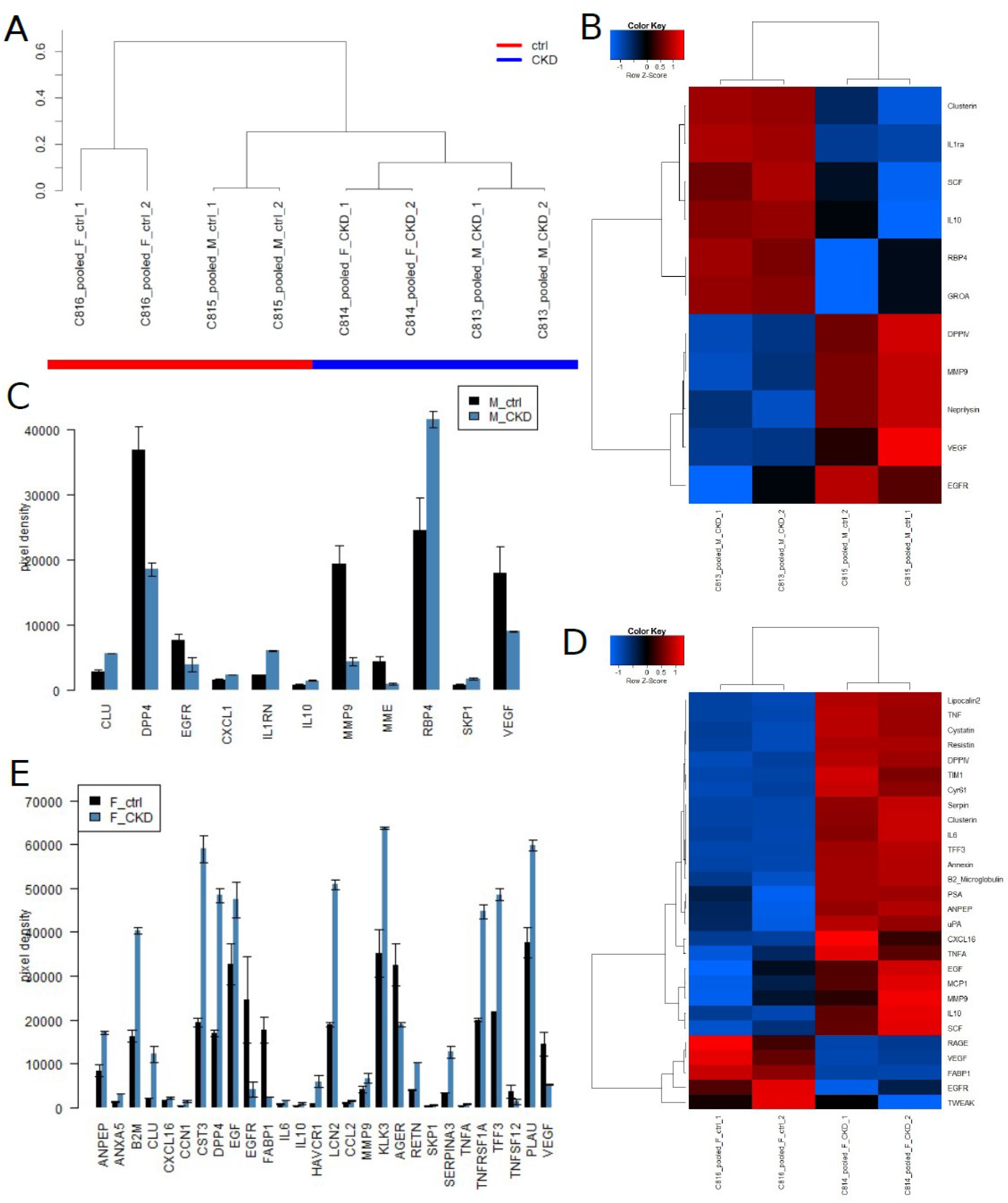
CKD biomarkers identified on the Human Kidney Biomarker Array in experiment series 1. (A) experiments cluster into CKD and healthy control based on the global kidney cytokine expression. (B) Heatmap and (C) barplot of markers in male (M) up-regulated (up): CLU, CXCL1, IL1RN, IL10, RBP4, SKP1, down-regulated (down): DPP4, EGFR, MME, MMP9, VEGF. (D) Heatmap and (E) barplot of markers in female (F) up-regulated (up): ANPEP, ANXA5, B2M, CCL2, CCN1, CLU, CXCL16, DPP4, EGF, IL6, IL10, HAVCR1, KLK3, LCN2, MMP9, PLAU, RETN, SERPINA3, SKP1, TNFA, TNFSRF14, TTF3, down-regulated (down): AGER, EGFR, FABP1, VEGF.

Figure 3 shows results of experiment series 2 on the Human Kidney Biomarker Array and the identified CKD biomarkers. On the global level of kidney cytokine expression the experiments clustered heterogeneously with respect to CKD and gender (Figure 3A). The heatmap (Figure 3B) and barplot (Figure 3C) show markers which were found up-regulated (up) in male (M): CLU, CXCL1, IL1RN, IL10, RBP4, SKP1, and down-regulated (down): DPP4, EGFR, MME, MMP9, VEGF. The heatmap (Figure 3D) and barplot (Figure 3E) show markers which were found up-regulated (up) in female (F): ANPEP, ANXA5, B2M, CCL2, CCN1, CLU, CXCL16, DPP4, EGF, IL6, IL10, HAVCR1, KLK3, LCN2, MMP9, PLAU, RETN, SERPINA3, SKP1, TNFA, TNFSRF14, TTF3, down-regulated (down): AGER, EGFR, FABP1, VEGF.

**Figure 3:**
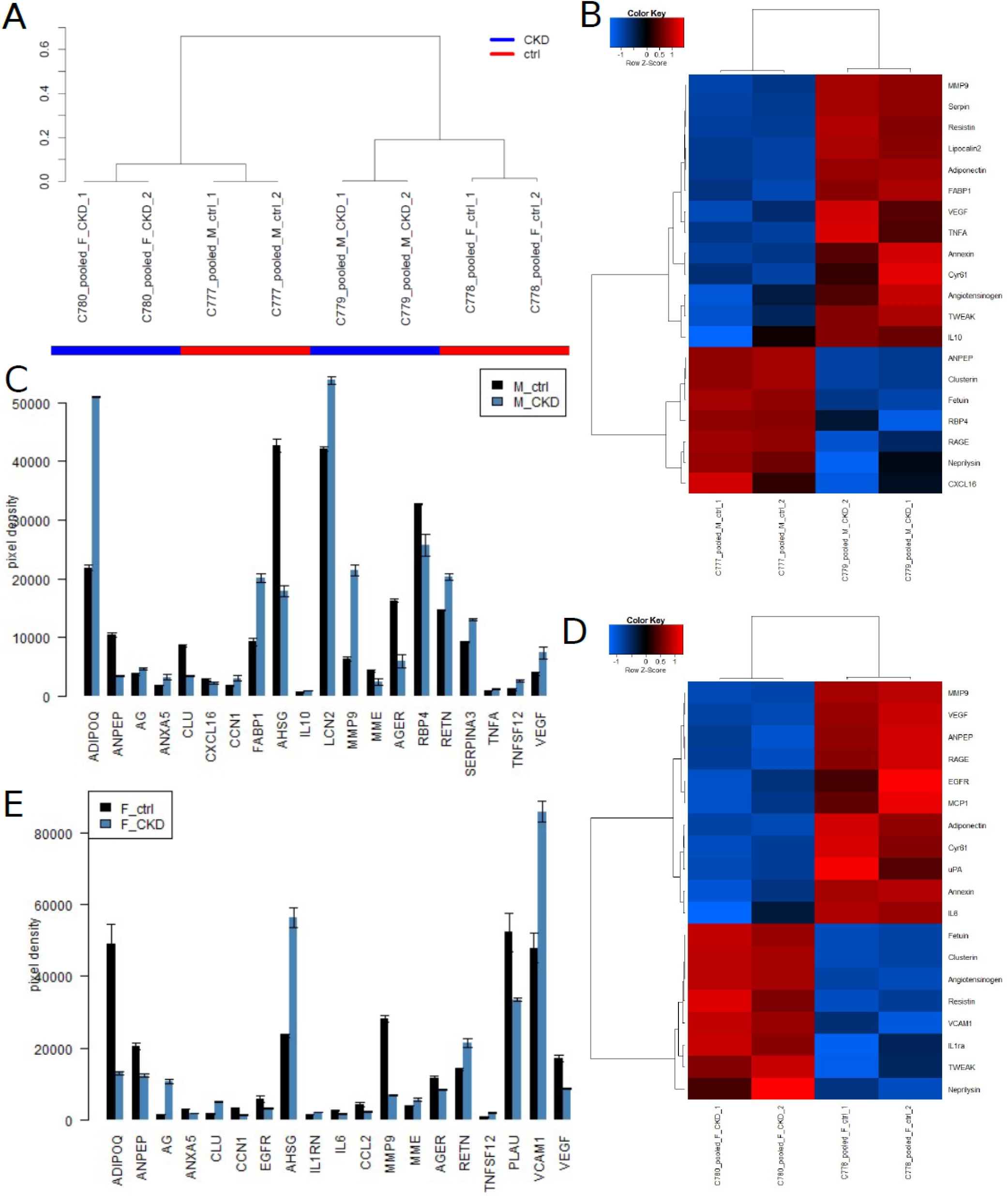
CKD biomarkers identified on the Human Kidney Biomarker Array in experiment series 2. (A) experiments cluster heterogeneously based on the global kidney cytokine expression. (B) Heatmap and (C) barplot of biomarkers in male (M) up-regulated (up): ADIPOQ, AG, ANXA5, CCN1, FABP1, IL10, LCN2, MMP9, RETN, SERPINA3, TNFA, TNFSF12, VEGF, down-regulated (down): AGER, AHSG, ANPEP, CLU, CXCL16, MME, RBP4. (D) Heatmap and (E) barplot of biomarkers in female (F) up-regulated (up): AG, AHSG, CLU, IL1RN, MME, RETN, TNFSF12, VCAM1, down-regulated (down): ADIPOQ, AGER, ANPEP, ANXA5, CCL2, CCN1, EGFR, IL6, MMP9, PLAU, VEGF.

### Summary of identified CKD biomarkers in experiment series 1 and 2

Table 2 shows a summary of CKD biomarkers identified in experiment series 1 and 2 for male CKD patients. Biomarkers identified in common in both series are underlined and in bold font. IL10 up-regulated and MME down-regulated in CKD are identified in common in both series.

**Table 1:**
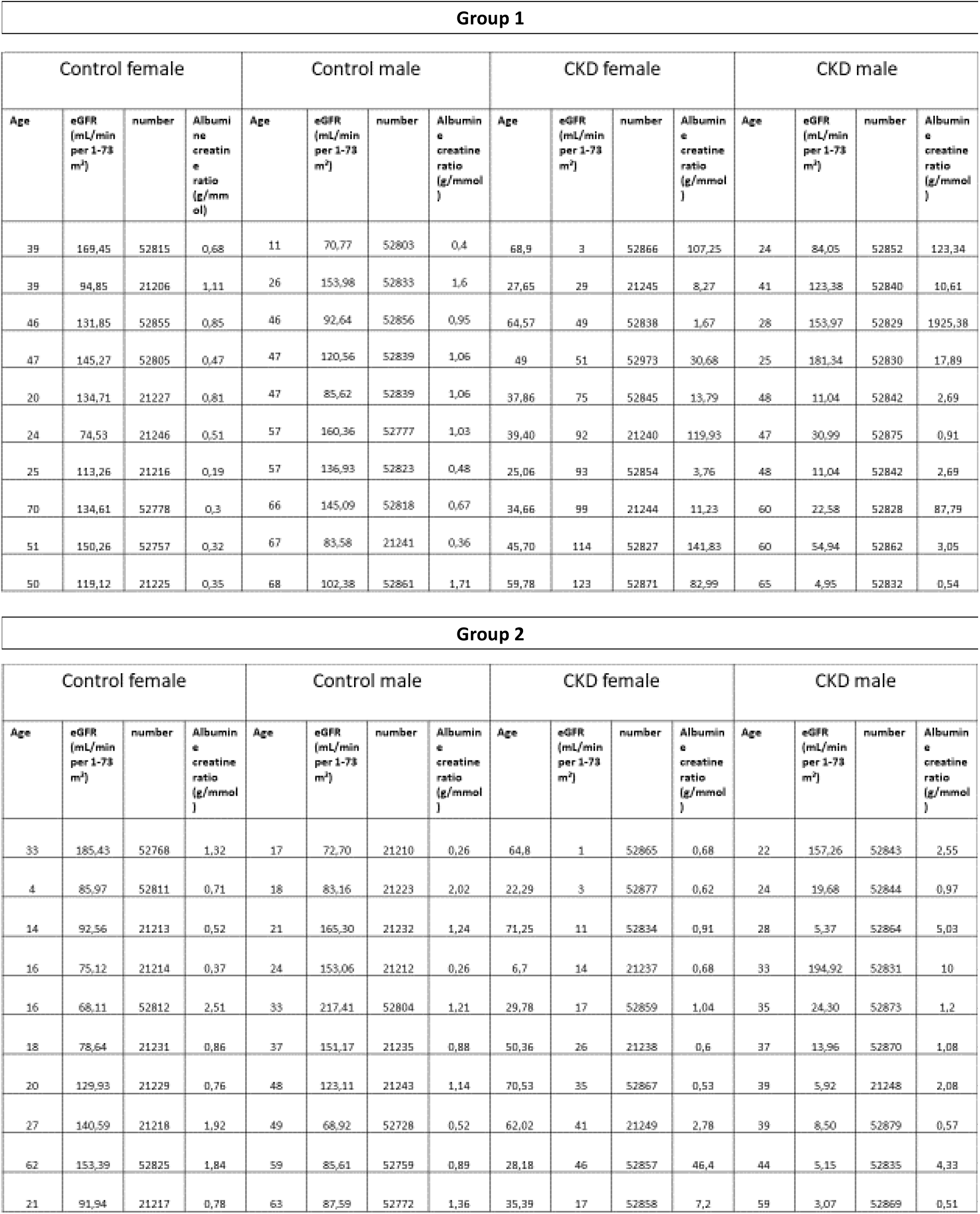
Characteristics of the patient cohort and the healthy individual cohort utilized in the cytokine assay analyses.

**Table 2:** CKD markers identified in male CKD patients in experiment series 1 and 2.

| cytokine | CKD_M_ExpSeries1 | CKD_M_ExpSeries2 |
| --- | --- | --- |
| ADIPOQ |  | up |
| AG |  | up |
| AGER |  | down |
| AHSG |  | down |
| ANPEP |  | down |
| ANXA5 |  | up |
| CCN1 |  | up |
| CLU | up | down |
| CXCL1 | up |  |
| CXCL16 |  | down |
| DPP4 | down |  |
| EGFR | down |  |
| FABP1 |  | up |
| IL1RN | up |  |
| **<u>IL10</u>** | **<u>up</u>** | **<u>up</u>** |
| LCN2 |  | up |
| **<u>MME</u>** | **<u>down</u>** | **<u>down</u>** |
| MMP9 | down | up |
| RBP4 | up | down |
| RETN |  | up |
| SERPINA3 |  | up |
| SKP1 | up |  |
| TNFA |  | up |
| TNFSF12 |  | up |
| VEGF | down | up |

Table 3 shows a summary of CKD biomarkers identified in experiment series 1 and 2 for female CKD patients. Biomarkers identified in common in both series are underlined and in bold format. CLU and RETN up-regulated and AGER, EGFR and VEGF down-regulated in CKD are identified in common in female in both series.

**Table 3:** CKD markers identified in female CKD patients in experiment series 1 and 2.

| cytokine | CKD_F_ExpSeries1 | CKD_F_ExpSeries2 |
| --- | --- | --- |
| ADIPOQ |  | down |
| AG |  | up |
| **<u>AGER</u>** | down | down |
| AHSG |  | up |
| ANPEP | up | down |
| ANXA5 | up | down |
| B2M | up |  |
| CCL2 | up | down |
| CCN1 | up | down |
| **<u>CLU</u>** | up | up |
| CXCL16 | up |  |
| DPP4 | up |  |
| EGF | up |  |
| **<u>EGFR</u>** | down | down |
| FABP1 | down |  |
| IL1RN |  | up |
| IL6 | up | down |
| IL10 | up |  |
| HAVCR1 | up |  |
| KLK3 | up |  |
| LCN2 | up |  |
| MME |  | up |
| MMP9 | up | down |
| PLAU | up | down |
| **<u>RETN</u>** | up | up |
| SERPINA3 | up |  |
| SKP1 | up |  |
| TNFA | up |  |
| TNFSF12 |  | up |
| TNFRSF14 | up |  |
| TTF3 | **up** |  |
| VCAM1 |  | **up** |
| **VEGF** | **down** | **down** |

### Identification of CKD biomarkers on the Human XL cytokine assay

After identification of CKD biomarkers on the Human Kidney Biomarker Array we set out to identify general cytokine markers without a prior association with the kidney as the ones on the kidney-specific cytokine assay. Figure 4 shows results of experiment series 3 on the human XL cytokine assay and the identified CKD inflammation-associated biomarkers. On the global level of cytokine expression the data segregated (Figure 4A) into two clusters of CKD (male and female) and healthy control (male). The heatmap (Figure 4B) and barplot (Figure 4C) show biomarkers which were found up-regulated (up) or down-regulated (down) in male (M). The heatmap (Figure 4D) and barplot (Figure 4E) show markers which were found up-regulated (up) or down-regulated in female (F). Interestingly, we found a large overlap between up-and down-regulated cytokines between both genders. Biomarkers in experiment series 3 overlapping between female (F) and male (M) up-regulated in CKD: APOA1 (up), ANGPT2 (up), C5 (up), CFD (up), GH1 (up), ICAM1 (up), IGFBP2 (up), IL8 (up), KLK4 (up), MMP9 (up), SPP1 (up). Markers in experiment series 3 overlapping between female (F) and male (M) down-regulated in CKD: FLT3LG (down), CSF1 (down), PDGFA (down), RETN (down), VEGFA (down).

**Figure 4:**
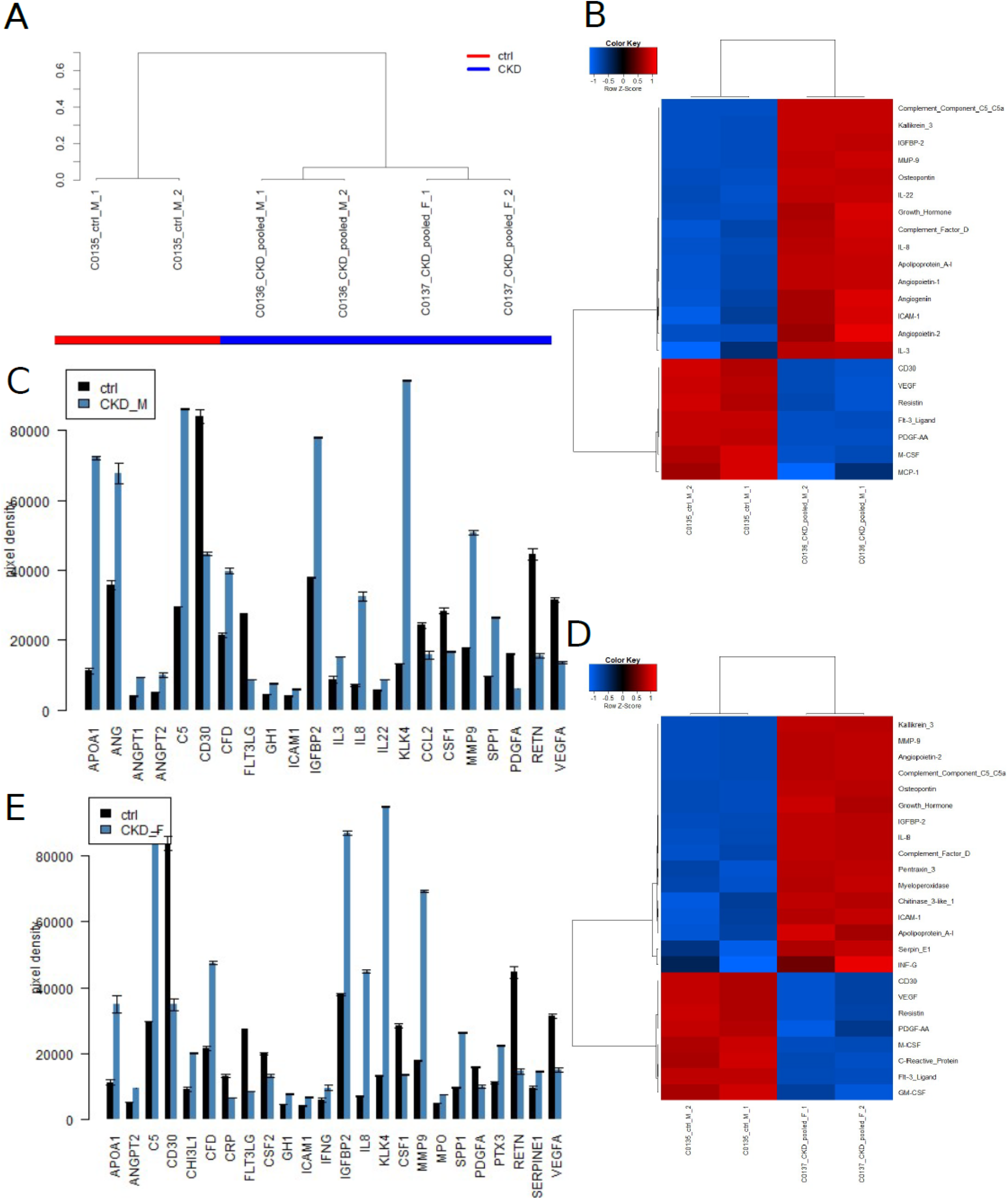
CKD biomarkers identified on the Human XL cytokine assay in experiment series 3 overlap in large parts between male and female. (A) data cluster into CKD and healthy control based on the global cytokine expression. (B) Heatmap and (C) barplot of markers in male (M) up-regulated (up) or down-regulated (down). (D) Heatmap and (E) barplot of markers in female (F) up-regulated (up) or down-regulated (down). Markers in experiment series 3 overlapping between female (F) and male (M) up-regulated in CKD: APOA1 (up), ANGPT2 (up), C5 (up), CFD (up), GH1 (up), ICAM1 (up), IGFBP2 (up), IL8 (up), KLK4 (up), MMP9 (up), SPP1 (up). Markers in experiment series 3 overlapping between female (F) and male (M) down-regulated in CKD: FLT3LG (down), CSF1 (down), PDGFA (down), RETN (down), VEGFA (down).

Table 4 shows a summary of CKD markers identified in experiment series 3 for male and female CKD patients and independent of gender taking the mean of male and female values. Markers identified in common in male and female are underlined and in bold format. This summary table highlights the large overlap between male and female CKD markers on the human XL cytokine platform.

**Table 4:** CKD biomarkers identified in the mean of male and female and male and female gender-specific in experiment series 3.

| cytokine | CKD_mean_M_F_ExpSeries3 | CKD_M_ExpSeries3 | CKD_F_ExpSeries3 |
| --- | --- | --- | --- |
| **<u>APOA1</u>** | <u>up</u> | <u>up</u> | <u>up</u> |
| ANG |  | up |  |
| ANGPT1 |  | up |  |
| **<u>ANGPT2</u>** | <u>up</u> | <u>up</u> | <u>up</u> |
| **<u>C5</u>** | <u>up</u> | <u>up</u> | <u>up</u> |
| CD3 | down |  | down |
| CHI3L1 | up |  | up |
| **<u>CFD</u>** | <u>up</u> | <u>up</u> | <u>up</u> |
| CRP | down |  | down |
| **<u>FLT3LG</u>** | <u>down</u> | <u>down</u> | <u>down</u> |
| CSF2 | down |  | down |
| **<u>GH1</u>** | <u>up</u> | <u>up</u> | <u>up</u> |
| **<u>ICAM1</u>** | <u>up</u> | <u>up</u> | <u>up</u> |
| IFNG | up |  | up |
| **<u>IGFBP2</u>** | <u>up</u> | <u>up</u> | <u>up</u> |
| IL3 |  | up |  |
| **<u>IL8</u>** | <u>up</u> | <u>up</u> | <u>up</u> |
| IL22 |  | up |  |
| **<u>KLK4</u>** | **<u>up</u>** | **<u>up</u>** | **<u>up</u>** |
| CCL2 |  | **<u>down</u>** |  |
| **<u>CSF1</u>** | **<u>down</u>** | **<u>down</u>** | **<u>down</u>** |
| **<u>MMP9</u>** | **<u>up</u>** | **<u>up</u>** | **<u>up</u>** |
| MPO | **<u>up</u>** |  | **<u>up</u>** |
| **<u>SPP1</u>** | **<u>up</u>** | **<u>up</u>** | **<u>up</u>** |
| **<u>PDGFA</u>** | **<u>down</u>** | **<u>down</u>** | **<u>down</u>** |
| PTX3 | **<u>up</u>** |  | **<u>up</u>** |
| **<u>RETN</u>** | **<u>down</u>** | **<u>down</u>** | **<u>down</u>** |
| SERPINE1 | **<u>up</u>** |  | **<u>up</u>** |
| **<u>VEGFA</u>** | **<u>down</u>** | **<u>down</u>** | **<u>down</u>** |

### Protein interaction network

Based on the cytokines differentially regulated on the human XL cytokine platform we set out to identify and characterize a protein interaction network involved in the inflammatory pathophysiology of CKD. Figure 5A shows the Protein interaction network generated via the STRING online tool ^14^. Via the STRING tool enriched disease associations with *Leukostasis* and *Artery Disease* (Figure 5C) were determined and a table of enriched Gene ontology Biological Processes (Figure 5D, top 25 terms sorted by strength is shown). These pointed to the inflammatory response and included the highlighted term *Regulation of kidney development* (strength=1.8, false discovery rate=0.0024). The proteins associated with Regulation of kidney development (PDGFA, MMP9, VEGFA) are highlighted in the network in Figure 5B.

**Figure 5:**
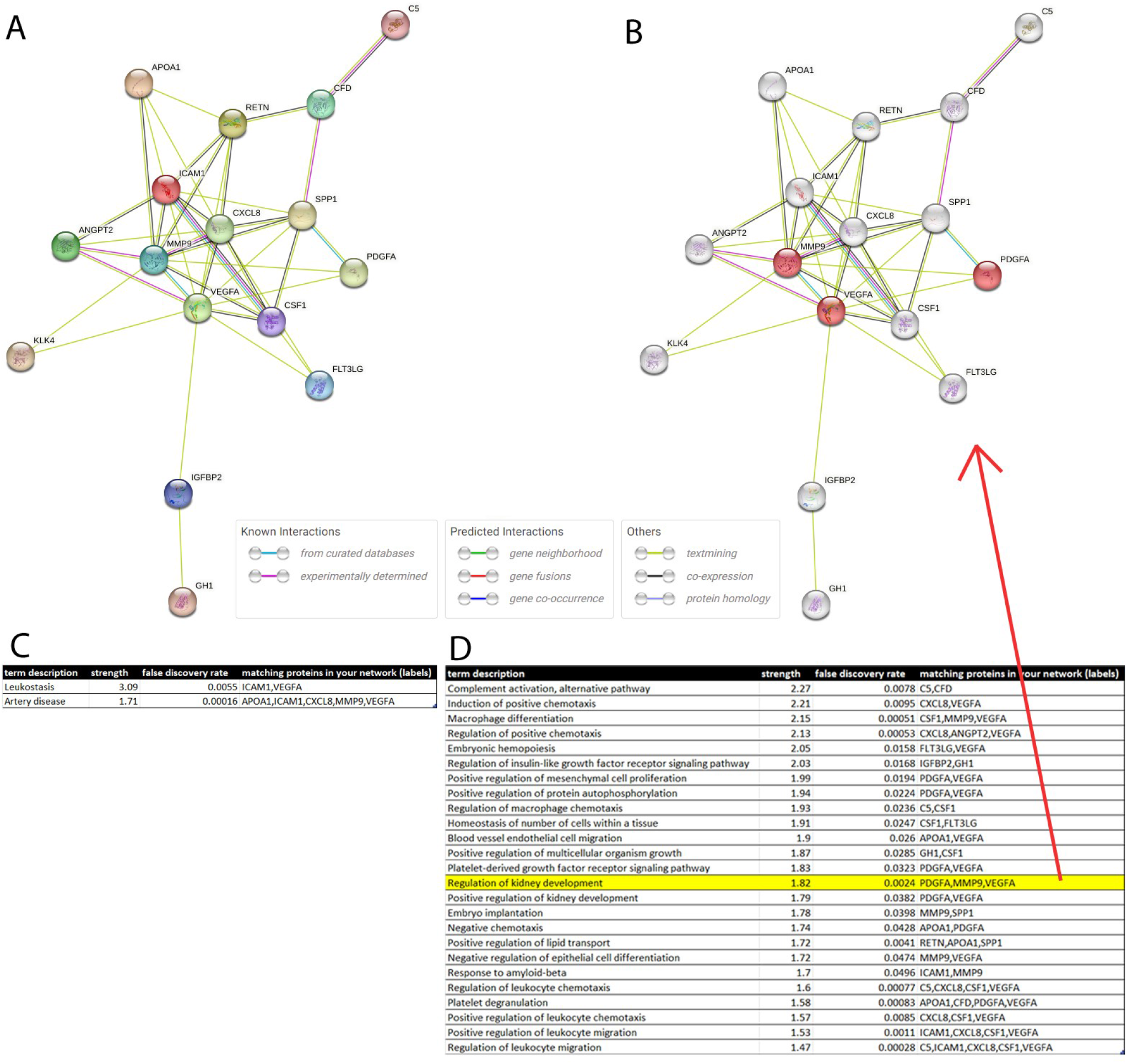
CKD biomarkers identified on the Human XL cytokine assay interact in a network regulating the inflammatory response affecting kidney development. (A) STRING protein interaction network (PPI) of CKD biomarkers identified on the Human XL cytokine assay. (B) PPI highlighting proteins involved in Regulation of kidney development. (C) Disease associations found enriched by the STRING tool are *Leukostasis* and *Artery Disease*. (D) Gene ontology Biological Processes found enriched point to the inflammatory response and include *Regulation of kidney development*.

## Discussion

In this two-phase analysis of cytokines in the urine of CKD patients and healthy individuals from Ghana, we identified urine-based cytokine biomarkers for CKD. In the first phase which was performed on a Human Kidney Biomarker Array we found variability between male and female and also between the two experiment series which consisted of pools of distinct individuals. This variability reflects the heterogeneity in CKD comprising a broad spectrum of distinct diseases such as diabetic nephropathy and focal segmental glomerulosclerosis. Furthermore, there is much diversity in the genetic background of CKD and recent findings on biomarkers associated with hitherto neglected areas such as telomeres, CNVs, mtDNA variants and sex chromosomes ^15^ may gain attention in the future. This may also help to explain the high heredity estimated at 30–75% which at the moment is incompletely understood ^15^. A further aspect of heterogeneity is found in the defintions of CKD which on the one hand have to cope with the complexity of the disease but on the other hand lead to strongly deviating asessments of CKD progression ^16^. Among the kidney-associated cytokines we found variable between genders and experiment series here LCN2 (alias NGAL; up-regulated in male in exp. Series 1 and in female in exp. Series 2, not-significant in the others) should be highlighted. NGAL had been established as marker of acute kidney injury but several problems including its unpredictable release have led to increasing concerns about its diagnostic value ^17^.

For the sake of robustness, only biomarkers regulated in the same direction in both experiment series are listed. The CKD biomarkers in male are: IL10 (Interleukin-10, up-regulated) and MME (Membrane metalloendopeptidase, down-regulated). Sinuani et al. describe that IL10 through increased proliferation of mesangial cells and mediated by several other cytokines induces progression of renal failure ^18^. Dedicated single nucleotide polymorphisms (SNPs) in the MME gene have been associated with a higher risk for diabetic nephropathy in female diabetes type 1 patients ^19^. The CKD biomarkers in female are: CLU (Clusterin) and RETN (Resistin, up-regulated) and AGER (advanced glycosylation end-product specific receptor, alias RAGE), EGFR (epidermal growth factor receptor) and VEGFA (vascular endothelial growth factor A, alias VEGF, down-regulated). CLU is described to be elevated in kidney disease ^20, 21^ although Guo *et al*. found that CLU deficiency worsens renal inflammation and tissue fibrosis after ischemia-reperfusion injury in mouse ^22^. For Resistin there are reports about elevated levels in CKD which are associated with decreased glomerular filtration rate and inflammation ^19^. AGER/RAGE was reported to be elevated in the serum of CKD patients ^23^. Up-regulation of EGFR has been described for CKD but EGFR inhibition in models of acute kidney injury (AKI) may also have deleterious effects ^24^.

In the second phase, we used a more general cytokine (human XL) assay and found a large percentage of cytokine biomarkers overlapping between both genders. The CKD biomarkers regulated in the same direction between male and female are: APOA1, ANGPT2, C5, CFD, GH1, ICAM1, IGFBP2, IL8, KLK4, MMP9 and SPP1 (up-regulated) and FLT3LG, CSF1, PDGFA, RETN and VEGFA (down-regulated). Interestingly, APOA1 is connected to APOL1 which has been associated with FSGS in haplotypes carried by patients of African descent. APOL1 is bound to HDL particles of which APOA1 is the major protein component but only 10% of APOA1-containing HDL particles have APOL1 ^25^.

We still need to confirm if indeed these APOL1-positive HDL particles play a special role in CKD. In comparison to the first phase where we saw strong gender-specific differences in the second phase we identified nearly the same markers regulated in the same direction for male and female. The large number of gender-specific differences found on the Human Kidney Biomarker Array in the first phase is in line with reports of gender differences in kidney function ^26^ and kidney disease ^27^. It is tempting to speculate that the more general cytokines measured on the human XL assay reflect more gender-independent inflammatory processes which preceed CKD.

Cytokines detected as differentially regulated on the Human XL cytokine assay as well as the Human Kidney Biomarker Array were RETN (Resistin, down-regulated on the Human XL cytokine assay), MMP9 (metalloproteinase 9, up-regulated on the Human XL cytokine assay) and VEGFA (down-regulated on the Human XL cytokine assay). VEGFA was regulated in the same direction on the Human Kidney Biomarker Array (down), MMP9 was up-in male in series 2 and female in series 1 and down-regulated in male in series 1 and female in series 2 but RETN in contrast to the XL assay was up-regulated. As mentioned above for RETN there are reports on elevated levels in CKD ^28^. MMP9 is described to be up-regulated in early stages of the disease and down-regulated in later stages ^29^. In our cohorts this might reflect the predominantly late disease stages in the distinct cohorts. For VEGFA there are conflicting data on the regulation in diabetic nephropathy going from up-regulation in rat models and also diabetic patients ^30^ over no effect ^31^ to down-regulation when diabetic nephropathy leads to glomerusclerosis ^30^. For glomerusclerosis & tubulorinterstitial fibrosis down-regulation has been reported ^30^.

We conclude, that our two-phase cytokine analysis of urine samples from CKD patients and healthy controls from Ghana revealed panels of kidney-specific cytokine biomarkers which were also gender-specific and a panel of gender-independent cytokine markers. The gender-specific markers are IL10 and MME for male and CLU, RETN, AGER, EGFR and VEGFA for female. The gender-independent cytokine markers were APOA1, ANGPT2, C5, CFD, GH1, ICAM1, IGFBP2, IL8, KLK4, MMP9 and SPP1 (up-regulated) and FLT3LG, CSF1, PDGFA, RETN and VEGFA (down-regulated).

## Methods

Participants in the present report were recruited from 2 academic medical centers in urban regions of Ghana between 2012– 2017. Persons with kidney disease were individuals aged 1-74 years with estimated glomerular filtration rate (eGFR) <60ml/min/1.73m^2^ (creatinine based chronic kidney disease epidemiology [CKD-EPI] collaboration equation without race adjustment)^1^ or albumin/creatinine ratio ≥3.0 mg/mmol (30 mg/g), and persons with a confirmed diagnosis of membranous glomerulonephritis, focal and segmental glomerulosclerosis/minimal change disease (FSGS/MCD), or childhood onset nephrotic syndrome. We excluded persons with obstructive uropathy, kidney tumors, multiple myeloma, polycystic kidney disease and women who were pregnant. Healthy persons without CKD were defined as individuals with eGFR ≥60ml/min/1.73m^2^ and albumin/creatinine ratio <3.0mg/mmol (<30 mg/g). Random urine samples were collected form cases and controls and aliquots of 10mls were taken into cryovials.

## Ethics statement

Ethics approval was obtained locally at each clinical site. The approval number for the Case-Control study is: GHSERC: 07/03/2013. Written informed consent was obtained from all participants.

Participants unable or unwilling to give consent or institutionalized were excluded.

### Cytokine assay experiments

The urine samples were analyzed using the Human Kidney Biomarker Array (ARY019) as well as the Human XL Cytokine Array kit (ARY022B) from Research And Diagnostic Systems, Inc. (Minneapolis, MN, USA), according to the manufacturer’s. For the first round of analysis the urine samples (control and CDK) were pooled prior to analysis. In the second round we analyzed another pool of urine samples (control and CDK).

In the third round urine from one individual male control sample was compared to the pooled male and female CKD samples. For each membrane 1 ml of urine was incubated overnight at 4°C on the respective membranes. Detection was carried out using a streptavidin-HRP and the ECL Prime Western-Blot-Detectionreagent (Merck, Darmstadt, Germany) detection reagents. The obtained signal were analyzed using FIJI / ImageJ software ^32^.

### Image analysis of cytokine assays

Scanned images of the cytokine assays were read into the FIJI / ImageJ software ^32^. The semi-automatic grid finding was based on pre-processing via Gaussian blur (size 4) and local maxima finding as described in Steinfath *et al*. ^33^. Local maxima were detected via the FIJI function “Find Maxima” and exported to a file in the csv format. The csv file containing the local maxima was imported into the R programming environment to detect the corners and interpolate the grid with the predefined size between the corners. The interpolation routine took into account alleyways between blocks within the grid and adjusted grid positions to local maxima where existent. For positions without detected local maxima the interpolated values were used. Grid positions were read into the FIJI Microarray Profile plugin by Bob Dougherty and Wayne Rasband (https://www.optinav.info/MicroArray_Profile.htm) which was employed to quantify the integrated densities of the spots. Grid positions were annotated with the cytokine identifiers provided in the manuals of the manufacturer (Proteome Profiler Array from R&D Systems, Human XL Cytokine Array Kit, Catalog Number ARY022B and Human Kidney Biomarker Array Kit, Catalog Number ARY019).

### Data analysis of cytokine assays

Integrated densities of the spots quantified by the FIJI Microarray plugin were imported into the R/Bioconductor ^34^ environment for further processing. Data was normalized via the Robust Spline Normalization from the R/Bioconductor package lumi ^35^. Cytokines were considered expressed when their integrated density was significantly (p < 0.05) over the background spots which were determined as spots with the minimum density plus 0.05 (max_density – min_density). Differential expression was assessed via the moderated t-test from the Bioconductor *limma* package ^36^. Limma p-values were adjusted for the false discovery rate (FDR) by the mehod from Benjamini and Hochberg ^37^. Cytokines were considered as differentially expressed via the criteria: detection p-value < 0.05 in at least one condition, ratio < 0.667 (down-regulated), ratio > 1.5 (up-regulated), limma-p-value < 0.05, FDR < 0.25. Heatmaps were generated with the function *heatmap*.*2* from the *gplots* package ^38^ using Pearson correlation as similarity measure. Bar plots were generated with the R-builtin function *barplot*. Dendrograms were produced via the R package *dendextend* ^39^ using Pearson correlation as similarity measure and complete linkage as agglomeration method.

## Author contributions

JA and VB conceived the study. BV, DA, TK, and EK recruited the patients and healthy individuals and collected the urine samples.

WW analysed the data and wrote the manuscript. LE and CT performed the cytokine assays and co-wrote the manuscript. JA supervised the work, co-wrote the manuscript and gave the final approval.

## Competing interests

The authors declare no competing interests.

## Acknowledgments

James Adjaye acknowledges support from the Medical faculty of Heinrich-Heine University, Duesseldorf.

Vincent Boima acknowledges support from the University of Ghana Medical School and all participants who consented to participate in this study.

## Data availability

Datasets for the analysis of cytokine assays were generated during the current study. The datasets are available in the supplementary data of this work.

## Supplementary Material

**Supplementary Table 1 (tableS1.xlsx): mean values and statistical test results of male patients CKD vs. healthy control in experiment series 1 on kidney cytokine array**.

**Supplementary Table 2 (tableS2.xlsx): mean values and statistical test results of female patients CKD vs. healthy control in experiment series 1 on kidney cytokine array**.

**Supplementary Table 3 (tableS3.xlsx): mean values and statistical test results of male patients CKD vs. healthy control in experiment series 2 on kidney cytokine array**.

**Supplementary Table 4 (tableS4.xlsx): mean values and statistical test results of female patients CKD vs. healthy control in experiment series 2 on kidney cytokine array**.

**Supplementary Table 5 (tableS5.xlsx): mean values and statistical test results of male patients CKD vs. healthy control in experiment series 3 on Human XL cytokine array**.

**Supplementary Table 6 (tableS6.xlsx): mean values and statistical test results of female patients CKD vs. healthy control in experiment series 3 on Human XL cytokine array**.

**Supplementary Table 7 (tableS7.xlsx): statistical test results of mean of male and female patients CKD vs. healthy control in experiment series 3 on Human XL cytokine array**.

**Supplementary Figure 1 (figS1.pptx): Scanned kidney cytokine arrays of experiment series 1.**

**Supplementary Figure 2 (figS2.pptx): Scanned kidney cytokine arrays of experiment series 2.**

**Supplementary Figure 3 (figS3.pptx): Scanned kidney cytokine arrays of experiment series 3**.

## Notes

### Competing Interest Statement

The authors have declared no competing interest.

