## Supplementary figures and images for "Urine-based detection of biomarkers indicative of chronic kidney disease in a patient cohort from Ghana"

### Supplementary Figure 1

#### Slide 1
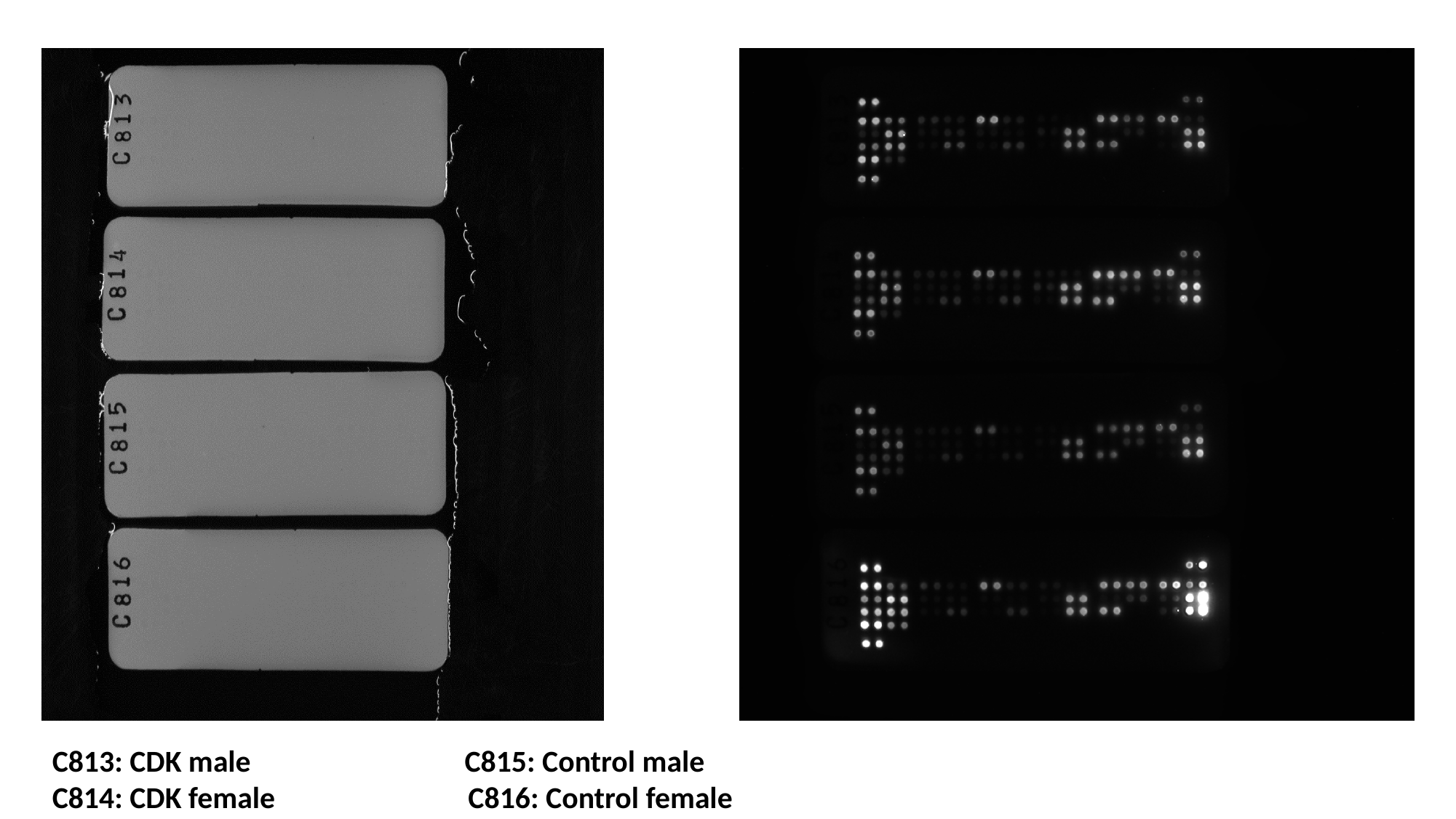

C813: CDK male C815: Control male
C814: CDK female C816: Control female

### Supplementary Figure 2

#### Slide 1
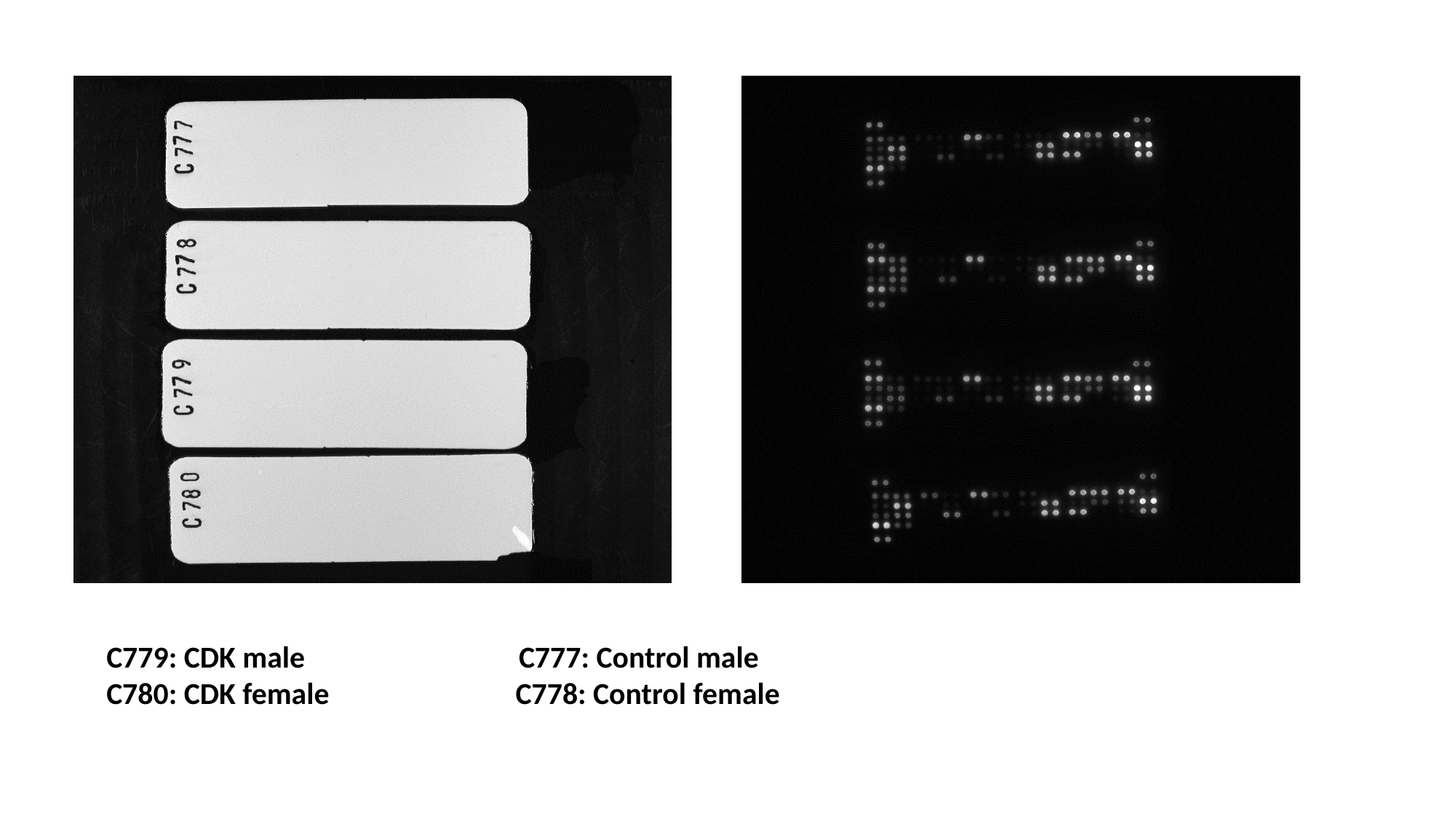

C779: CDK male C777: Control male
C780: CDK female C778: Control female

### Supplementary Figure 3

#### Slide 1
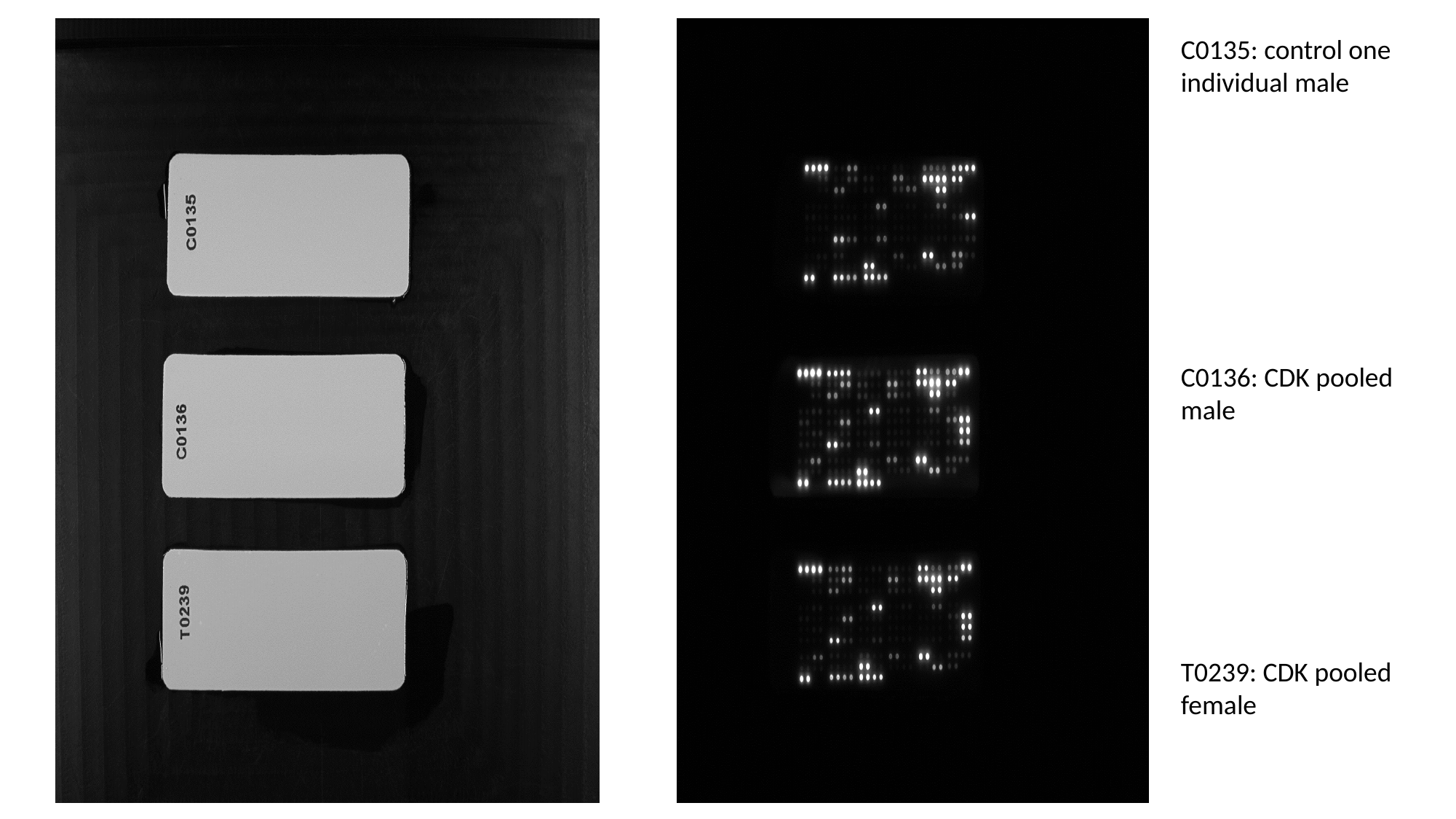

C0135: control one individual male
C0136: CDK pooled male
T0239: CDK pooled female
